# The Missing Link: Predicting Connectomes from Noisy and Partially Observed Tract Tracing Data

**DOI:** 10.1101/063867

**Authors:** Max Hinne, Annet Meijers, Rembrandt Bakker, Paul H. E. Tiesinga, Morten Mørup, Marcel A.J. van Gerven

## Abstract

Our understanding of the wiring map of the brain, known as the *connectome*, has increased greatly in the last decade, mostly due to technological advancements in neuroimaging techniques and improvements in computational tools to interpret the vast amount of available data. Despite this, with the exception of the *C. elegans* roundworm, no definitive connectome has been established for any species. In order to obtain this, tracer studies are particularly appealing, as these have proven highly reliable. The downside of tract tracing is that it is costly to perform, and can only be applied *ex vivo*. In this paper, we suggest that instead of probing all possible connections, hitherto unknown connections may be predicted from the data that is already available. Our approach uses a ‘latent space model’ that embeds the connectivity in an abstract physical space. Regions that are close in the latent space have a high chance of being connected, while regions far apart are most likely disconnected in the connectome. After learning the latent embedding from the connections that we did observe, the latent space allows us to predict connections that have not been probed previously. We apply the methodology to two connectivity data sets of the macaque and we demonstrate that the latent space model is successful in predicting unobserved connectivity, outperforming two alternative baselines in nearly all cases. Furthermore, we show how the latent spatial embedding may be used to integrate multimodal observations (i.e. anterograde and retrograde tracers) for the mouse neocortex. Finally, our probabilistic approach enables us to make explicit which connections are easy to predict and which prove difficult, allowing for informed follow-up studies.

## 1. Introduction

Recent years have seen a surge in research effort devoted to obtaining the human *connectome*, a map of all the connections in the human brain at the level of macrosopic brain regions (Sporns et al. 2005, Hagmann 2005). Technological advances, in particular diffusion-weighted MRI (dMRI), have enabled bundles of white-matter fibers to be identified *in vivo* in unprecendented detail. However, dMRI suffers from a number of drawbacks (Jones et al. 2013, Schultz et al. 2014, Reveley et al. 2015). For instance, it is an indirect measuring technique (Jbabdi et al. 2015): Rather than directly observing axons or large fiber bundles, these must be inferred from the diffuse movement of water molecules, using a process called tractography. In practice, the problem tractography tries to solve may be underspecified, as a single voxel may contain fibers that cross, ‘kiss’, merge or split (Jbabdi & Johansen-Berg 2011). As a result, it may be unclear which path the estimated fibers follow. Further problems arise when interpreting the output of (probabilistic) tractography. The number of streamlines (i.e. candidate fiber trajectories) that connect two regions of interest is often used synonymously with fiber count, yet the actual number of streamlines between two regions is an intricate function of the actual fiber count and several parameters of the dMRI acquisition and tractography procedure (Jones et al. 2013, O’Donnell & Pasternak 2015). In all, despite dMRI having greatly advanced the field of connectomics by being applicable in living human subjects, it is far from the be-all end-all solution to finding gross anatomical connectivity.

Earlier approaches for studying brain connectivity (Catani et al. 2013) involve techniques such as post-mortem dissection (Broca 1861, Gall & Spurzheim 1812) as well as tract tracing in animal subjects (Köbbert et al. 2000). In the latter approach a tracer (such as a fluorescent dye or a virus) is injected into neuronal tissue of a living animal. After appropriate waiting time, the animal is sacrificed to allow the tracer material to spread through the tissue, either in the direction from cell soma to axon terminal (known as anterograde tracing), or vice versa (retrograde tracing). Inspection of the virus expression or the fluorescence of the dye is subsequently used to determine to which other neurononal populations the injection site was connected (Oztas 2003, Lanciego & Wouterlood 2011, Lu 2011). Tract tracing has a number of advantages over dMRI-based connectivity estimation. First of all, tract tracing provides unequivocal proof that two regions are connected. In dMRI, there is always a possibility that fiber tracts follow the same highway, but do not mix. Furthermore, tract tracing can recover the direction of the tracts it recovers, something which is impossible to do with dMRI. Furthermore, the probed connections are measured directly, without the need for an additional processing step such as tractography. This results in very accurate connectivity estimates, in particular regarding long-range connections (Dauguet et al. 2007, Jbabdi et al. 2015) and has prompted researchers to use tract tracing methods as a means to evaluate the performance of dMRI-based structural connectivity estimation (Azadbakht et al. 2015, Dyrby et al. 2007, Seehaus et al. 2015).

Compared to dMRI, tract tracing is a very expensive procedure for probing connectivity (Bakker et al. 2012, Sporns 2010). It requires sacrificing animal subjects, as well as substantial manual labor in administering the tracers and processing the treated brain tissue. Through a process known as ‘link prediction’ (Lü & Zhou 2011, Liben-Nowell & Kleinberg 2007, Clauset et al. 2008) the number of experimental studies needed to evaluate all possible connections in a connectome may be reduced. The general idea behind this technique is that the connections that have been observed carry enough information for the missing connections to be predicted. One class of models used for making link predictions assumes that connections are the result of hidden properties of the nodes in the network (i.e. regions of interest or neuronal populations). For instance, stochastic block models assume the nodes of a network have latent class labels, and that the probability of a connection between two nodes depends on whether they share the same label (Nowicki & Snijders 2001). By learning this latent structure from the data, i.e. which node has which label, new connections (or the absence thereof) may be predicted (Guimerà & Sales-Pardo 2009, Kemp et al. 2006, Herlau et al. 2014, Mørup et al. 2010, Schmidt & Mørup 2013). The concept of latent node classes also forms the basis of community detection (Newman 2010), for which the goal is to identify sets of nodes that have more connections among themselves than with nodes outside the set. Another latent structure approach assumes that the nodes of a network are actually embedded in an unknown physical space (a ‘latent space’) (Bullock et al. 2010, Barthélemy 2011). When a latent space model (LSM) is used for link prediction, the (Euclidean) distance between the positions of nodes in the latent space is used to determine the likelihood of a connection. This approach is clearly applicable when networks represent geographically restricted phenomena, like traffic, power grids and the internet, but may also be used in more abstract settings, such as a social network with ties dependent on political ideology, rather than spatial location (Bullock et al. 2010). In this sense, LSM subsume latent class models, as one of the (non-spatial) dimensions of the model can simply reflect class label, making such models a special case of LSM (Miller et al. 2009).

While stochastic block models have been used for modeling and prediction of links in structural connectivity (Hinne et al. 2015, Ambrosen et al. 2013, 2014), LSMs have so far mostly been applied to social network analysis (Hoff et al. 2002, Sarkar & Moore 2005, Sewell & Chen 2015) instead of to structural connectivity. However, clearly the connectome is spatially embedded (Zitin et al. 2014, Bullmore & Sporns 2012, Ercsey-Ravasz et al. 2013), suggesting that the use of LSM can improve the quality of link prediction. In the current study, we describe an extended probabilistic LSM with which we embed tract-tracing connectomes into a latent space. This allows us to predict unknown connections in macaque visual cortex (Felleman & Van Essen 1991) and macaque cerebral cortex (Markov et al. 2014). Additionally, the procedure is applied to combine anterograde and retrograde tract tracing data for the mouse neocortex (Zingg et al. 2014). While in this data set all connections have been observed, the different tracer directions disagree about connection strengths. We show that by embedding the network into a latent space, both sources of data can be explained by a single connectome. The probabilistic nature of our approach provides an intuitive representation of the uncertainty in the parameters we estimate, in the form of their posterior distribution. This uncertainty may be used to determine which predicted connections are reliable and which require more data to be estimated with confidence.

**Table 1.**
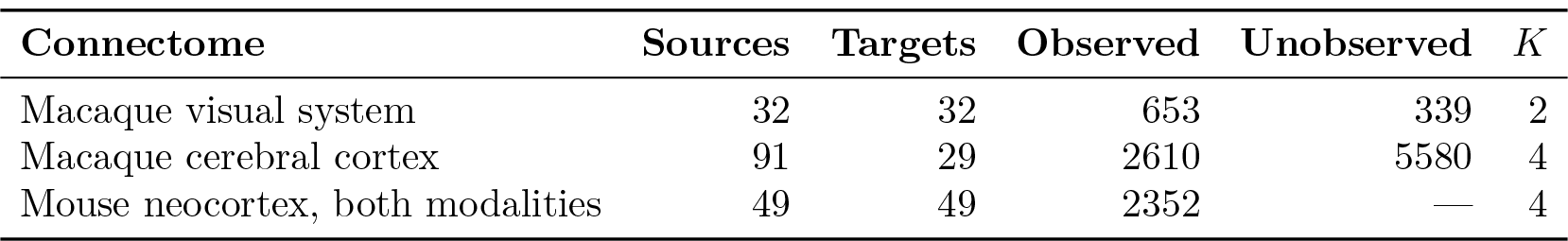
For each of the different connectivity data sets, the table shows the number of source nodes, the number of target nodes, the numbers of observed and unobserved connections and finally the number of observed connection strength classes *K*.

| Connectome | Sources | Targets | Observed | Unobserved | $K$ |
| --- | --- | --- | --- | --- | --- |
| Macaque visual system | 32 | 32 | 653 | 339 | 2 |
| Macaque cerebral cortex | 91 | 29 | 2610 | 5580 | 4 |
| Mouse neocortex, both modalities | 49 | 49 | 2352 | — | 4 |

## 2. Materials and Methods

### 2.1. Data

The data sets used in this paper are publicly available. Surface data was available for the macaque data sets, but not for the mouse data as the node definitions for this data set are layer-specific. The properties of each of the data sets are summarized in Table 1 and discussed in detail below.

#### 2.1.1. Macaque visual system

The macaque visual system connectome consists of the combined results of 31 studies, collected by Felleman & Van Essen (1991). The result is a partially observed connectome of size 32 × 32, consisting of both anterograde and retrograde tracings for one hemisphere. Connections are classified as either absent, present or unknown. Of the 32 · 31 = 992 possible connections, 653 candidate connections have been probed and of these, 286 are considered to represent connected node pairs. The other 339 connections remain unknown, and will be predicted using the proposed method.

#### 2.1.2. Macaque cerebral cortex

A macaque cerebral cortex connectome was obtained by Markov et al. (2014) by injecting retrograde tracers into 29 of 91 architectonic areas, all mapped to the left hemisphere. The result is a partially observed connectome of size 91 × 29. Connection strengths are quantified using the extrinsic fraction of labeled neurons (FLNe) index, which is the fraction of labeled neurons in the source area (i.e. those that send a projection to the injection site), divided by the total number of labeled neurons in the brain except for those in the injection area. Although these scores provide a continuous scale, Markov et al. (2014) propose a set of thresholds to categorize the connections into strong, moderate, sparse and absent. Throughout this paper, we use this ordinal representation to predict the unobserved connections.

*Mouse neocortex*. Zingg et al. (2014) have collected both anterograde and retrograde tracings for the mouse neocortex, which have been aggregated into two 49 × 49 connectivity matrices, shown in Fig. 1A–B, for which the connection strengths have been manually assigned to the categories strong, moderate, sparse and absent. While the connectomes have been fully observed and therefore contain no missing connections, the anterograde and retrograde tracings are not in complete agreement, as is shown in Fig. 1C. This may for example be due to experimental variability, e.g. differences in volume or location of injections) or different sensitivity for the retrograde or anterograde tracers. It is unclear how the two data sources may best be combined. In (Zingg et al. 2014), the combination is performed using a logical AND-operator on the two matrices: a connection is considered present if it is present in both observations. The result is a binary connectome, in which the information contained in the connection strengths is effectively lost. In the following, we will use our methodology to estimate a single connectome using both sources of data, thus cleaning up and reconciling the experimental variability.

### 2.2. The latent space model

The goal of our method is to predict connectivity for potential connections for which no tracer data is available, informed by the connections that do have observed data. To accomplish this, we assume that the *p* nodes of the network are in fact embedded in a latent space with dimensionality *D*, so that each node *i* has a latent position **z***_i_* ∈ ℝ*^D^* (Bullock et al. 2010, Hoff et al. 2002, Sarkar & Moore 2005, Zitin et al. 2014, Barthélemy 2011). Furthermore, we assume that the propensity for two nodes to be connected, or the strength of such a connection, depends on the distance *l*_*ij*_ = ‖**z**_*i*_ – **z**_*j*_‖_2_ between the two nodes in the latent space. If no tracer data was available, the nodes are considered to be distributed uniformly within this latent space. As soon as connections between pairs of nodes become observed, this latent arrangement becomes constrained — for example, nodes that are strongly connected should be close to each other and conversely, disconnected nodes should be far apart. The higher the dimensionality of the latent space, the more complex configurations of the connectome the model can represent. For example, in a 1D model the latent positions are ordered on a line, which can host only a limited number of different connectivity structures. On the other hand, in a high-dimensional space the degrees of freedom of the model will be sufficiently high to capture a more complex network topology (although for our purposes, a high-dimensional latent space will be prone to overfitting).

**Figure 1.**
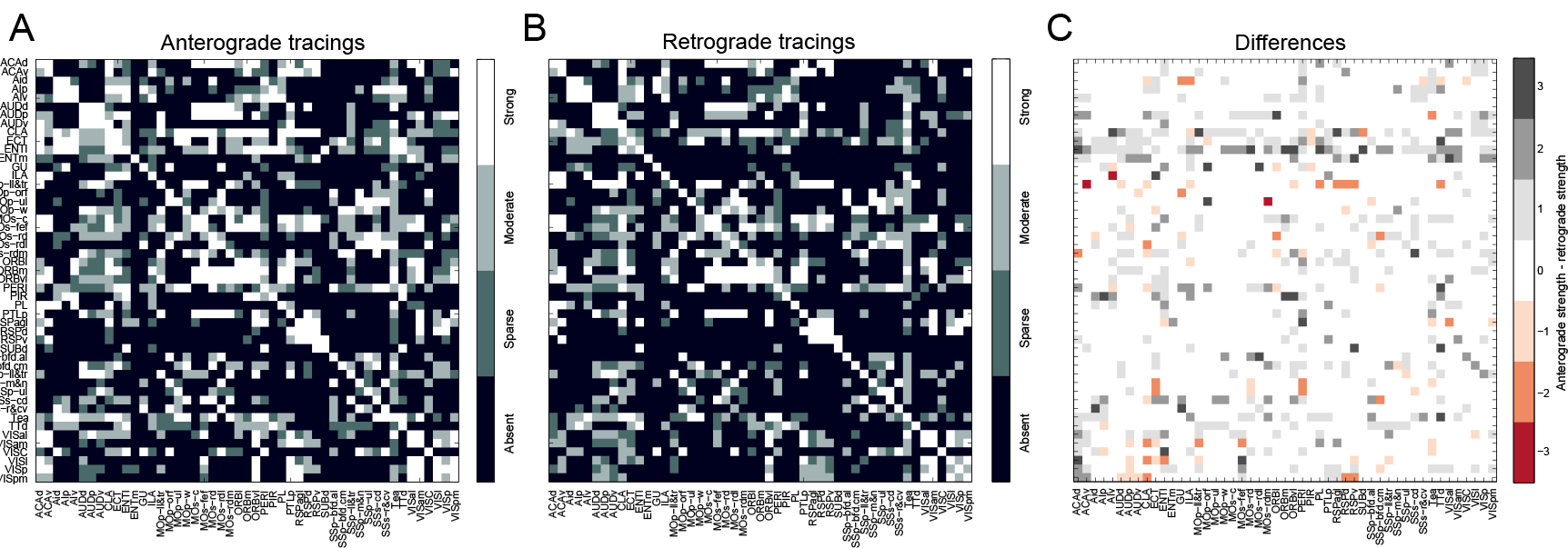
The mouse neocortex data (Zingg et al. 2014) **A** Shows the anterograde tracing result, **B** shows the retrograde tracing result, and **C** shows the differences between the two, using the following numerical representation for connection strengths: 0: absent, 1: sparse, 2: moderate and 3: strong.

As tracer data is typically available in a thresholded form, e.g. binary connections or ordinal connection weights, the latent space is accompanied by a set of boundaries that determine which range of distances corresponds to a particular connection weight. This idea is implemented using an ordinal regression model (Hoff 2008, Chu & Ghahramani 2005). It defines the probability of an ordinal connection class *k* between nodes *i* and *j* as *f*_*ijk*_ = Φ (*i*, *j*, *k*) – Φ(*i*, *j*, *k* − 1), in which

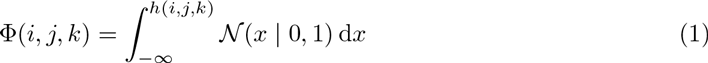

gives the cumulative density of the standard normal distribution on the interval [−∞, *h*(*i*, *j*, *k*)]. Here, 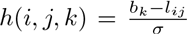 serves to scale and translate the Euclidean distance in the latent space to the intervals of the normal density function. Note that *b*_*k*_ and σ are the same for all connections. The categorical observed connection weights **A** = {*a*_*ij*_} are assumed to follow a categorical distribution with probability vector **f**_*ij*_, subject to 0 ≤ *f*_*ijk*_ ≤ 1 and ∑_*k*_ *f*_*ijk*_ = 1. If anterograde and retrograde tracer data are available separately, as in the data collected by Zingg et al. (2014), both **A** = {*a*_*ij*_} and **R** = {*r*_*i*j_} follow such a distribution.

Importantly, once the latent positions **z**_*i*_ and the class boundaries *h*_*ijk*_ have been learned using the available observations, the same parameters can be used to predict the class weight probabilities **f**_*ij*_ for unobserved connections, and subsequently predict **A**. Thus, the latent space model as described here serves as a mechanism to ‘complete’ a partially observed connectome.

The latent space model describes only symmetric connectivity behavior, as the Euclidean distance is a symmetric function. However, some nodes may be more prone to incoming or outgoing connections than distance alone can explain. For example, some nodes may be hubs — nodes which a large number of connections compared to the rest. To allow this phenomenon in our model, we add an ‘asymmetric effect’ which is captured in the vectors δ ∈ℝ^p^ and ε ∈ℝ^p^ that model the additional likelihood of nodes having incoming and outgoing connections, respectively (Wang &Wong 1987). The earlier distance measure is then replaced by

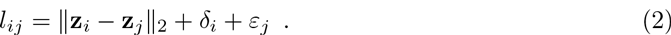

The generative model complete with priors on hyperparameters is shown in A, which also discusses constraints to make the model identifiable. Importantly, connections for which no tracer data is observed, provide no information to the model. Instead, by finding the spatial embedding of nodes within the latent space, the ordinal probabilities **f**_*ij*_ for these unknown connections may be inferred. The result is a probabilistic connectome that assigns for each connection a probability for each ordinal category.

To compute the posterior distribution of the latent locations and connection class probabilities, a Hamiltonian Monte Carlo sampling scheme is used as described in more detail in B. The result of this procedure is a collection of samples that collectively represent the distribution over the parameters of interest, i.e. the latent positions and the unobserved connection weights.

### 2.3. Optimal number of dimensions

To determine the optimal dimensionality of the latent space, 10-fold cross-validation was used for each of the different data sets. For each fold, the model was trained using nine-tenth of the observed connections and evaluated using the likelihood of the remaining one-tenth. To evaluate the performance of different numbers of dimensions, the parameter *D* was varied in the range [1,…, 6]. The dimensionality 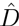 that resulted in the best generalizability (i.e. highest likelihood on the withheld data) was considered the optimal dimensionality. The model was then trained using 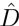 and all available data.

### 2.4. Prediction error and uncertainty

The performance of the predicted connectivity is evaluated using the cross-validation results. Per fold, we first compute for the *t*th collected sample 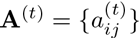 the absolute difference between the predicted connection weights 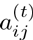 and the actual connection weight *a*_*ij*_, which is subsequently averaged over all connections, i.e.

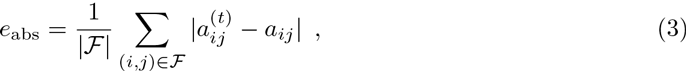

in which 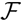 is the set of edges (*i*, *j*) in that particular cross-validation fold. This error measure is in the range [0, *K* – 1]. Note that *e*_abs_ is a conservative measure of the performance, as Bayesian averaging would typically reduce the error. However, to be consistent with the error measures described next, we evaluate *e*_abs_ per sample.

Depending on the intended application of the predictions, it may be more relevant to consider only the presence or absence of a connection instead of the difference in connection weight. In other words, predicting a weak connection that should have been absent may be a more severe error than predicting a strong connection that is in fact only moderate. We therefore also compute the false positive rate *e*_fpr_ and false negative rate *e*_fnr_, as

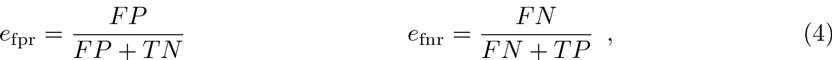

with the terms given by 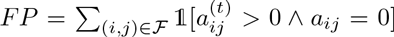, 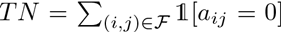, 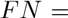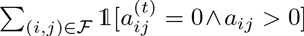 and 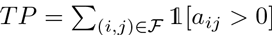. For each error measure, the presented results are subsequently averaged over all samples and over the cross-validation folds.

In addition to the prediction error, the probabilistic approach to latent space models allows us to compute the uncertainty that is associated with the predictions. To do so, we consider the posterior distributions over the parameters of interest. The width of these distributions provides a measure of uncertainty. If the model is certain about a prediction, the posterior will be peaked around the mode of the distribution, but if there is a lot of uncertainty the distribution will be more dispersed. To analyze how certain the predictions are, we quantify for each of the estimated parameters *f*_*ijk*_ the associated uncertainty as the 95% credible interval, i.e. the width of the range in which 95% of the posterior probability density lies. For each connection between node pairs (*i*, *j*), the largest uncertainty of the *K* possible connection strength classes is reported as the final measure of uncertainty.

### 2.5. Baseline predictions

Two baselines were constructed in order to interpret the results from the latent space model. In the first, connection probabilities for a particular connection strength class *k* are determined by the fraction of connections in the training data having connection weight *k*. In this baseline, the probability vector **f**_*ij*_ is the same for all pairs (*i*, *j*). This approach corresponds to a naive data imputation method, and its performance demonstrates how much of the connectivity prediction can be done by the connection weight distributions alone. To compute false positive and false negative rates (see previous section) in a similar fashion to the LSM, a posterior distribution is constructed for the baseline by drawing samples. In each sample, each connection 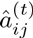 is drawn from a Bernoulli distribution with the probability as just described. The sampled connections are then treated the same as for the LSM approach.

In the second baseline, a zero-dimensional latent space is used. In other words, the only flexibility the model has, is in the directional effects δ and *ε*. This baseline serves to evaluate the additional effect of the latent space, compared to the predictive performance of the degree distribution of the training data.

### 2.6. Relative degree and clustering

The predicted connectivity can be analyzed in a number of ways. First, we present the expected posterior connectome, which serves as a summary of the predictions. It is given by **Â** = {â_*ij*_}, with

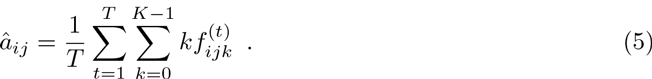

Using the posterior expectation, we then compute for each node *i* its normalized observed degree, i.e. 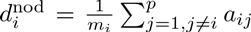 in which *m*_*i*_ indicates the number of observed connections for node *i* and its normalized predicted degree 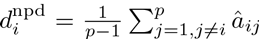, which considers all possible connections.

In addition, we consider clustering of the predicted connectome. Here, the weighted and directed network Â serves as input for a network clustering procedure from the Brain Connectivity Toolbox (Rubinov & Sporns 2010). The quality of the clustering is expressed using the modularity measure (Leicht & Newman 2008), given as

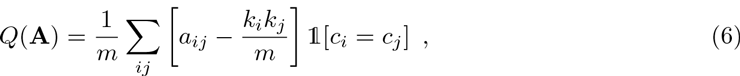

in which *m* is the sum of all connection weights, *k*_*i*_ and *k*_*j*_ are the sum of weights connected to nodes *i* and *j* and finally *c*_*i*_ and *c*_*j*_ represent the cluster labels of nodes *i* and *j*, respectively. Modularity reflects the number of connections found within clusters, compared to chance level, so that higher values indicate that there are many within-cluster connections and few between-cluster. For comparison, we also determine the clustering for the predictions using the baselines described in the previous section. However, the second baseline did not result in sensible clusterings (as concluded from a low modularity scores and a large number of very small clusters), which is why these have been omitted from discussion in the remainder.

## 3. Results

The latent space model is applied to each of the data sets described in Section 2.1. For the first two data sets, the macaque visual system tracer data and the macaque cerebral cortex tracer data, the goal is to predict the unobserved connections, i.e. to complete the partially observed connectome. For the third data set, the mouse neocortex, all connections have been observed, but here the latent space model serves to unite the anterograde and retrograde tracer data, i.e. to identify the connectome from which both data modalities originate. In either setting, the optimal dimensionality of the latent space must first be determined, using the cross-validation approach as described in Section 2.3. The resulting optimal number of dimensions 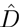 are shown in Table 2. In C, the generalization performance is shown for each of the considered dimensionalities, together with the corresponding posterior expectations.

**Table 2.** For each of the different data sets, the table shows the optimal latent dimensionality 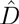 the prediction errors using the baseline models and the LSM, and finally the prediction uncertainty of the LSM. Standard deviations are indicated within parentheses, where applicable. The error for the mouse neocortex using both data modalities is the average error of the predictions with either data source. The different error measures are absolute error *e*_abs_ (top row per connectome), false positive rate *e*_fpr_ (middle row per connectome) and false negative rate *e*_fnr_ (bottom row per connectome). Per combination of connectome and error measure, the best performance is indicated in boldface.

| Connectome | $\hat{D}$ | Error | | | Uncertainty |
| --- | --- | --- | --- | --- | --- |
|  |  | Baseline 1 | Baseline 2 | LSM |  |
| Macaque visual system | 2 | 0.47 (0.03) | 0.40 (0.01) | **0.12 (0.03)** | 0.36 (0.29) |
|  |  | 0.22 (0.00) | 0.26 (0.01) | **0.10 (0.02)** |  |
|  |  | 0.42 (0.00) | 0.31 (0.02) | **0.12 (0.04)** |  |
| Macaque cerebral cortex | 1 | 1.22 (0.05) | 1.04 (0.01) | **0.76 (0.02)** | 0.37 (0.01) |
|  |  | 0.38 (0.00) | 0.34 (0.01) | **0.27 (0.01)** |  |
|  |  | 0.28 (0.00) | 0.24 (0.00) | **0.18 (0.01)** |  |
| Mouse neocortex, anterograde | 2 | 1.41 (0.04) | 1.02 (0.04) | **0.61 (0.04)** | 0.36 (0.20) |
|  |  | 0.29 (0.00) | 0.25 (0.01) | **0.16 (0.02)** |  |
|  |  | 0.37 (0.00) | 0.33 (0.01) | **0.22 (0.02)** |  |
| Mouse neocortex, retrograde | 2 | 1.38 (0.05) | 0.99 (0.04) | **0.64 (0.04)** | 0.34 (0.21) |
|  |  | 0.25 (0.00) | 0.26 (0.01) | **0.19 (0.01)** |  |
|  |  | 0.40 (0.00) | 0.33 (0.01) | **0.20 (0.01)** |  |
| Mouse neocortex, both modalities | 4 | 1.40 (0.04) | 0.96 (0.04) | **0.55 (0.04)** | 0.31 (0.20) |
|  |  | 0.27 (0.00) | 0.24 (0.01) | **0.14 (0.01)** |  |
|  |  | 0.39 (0.00) | 0.35 (0.00) | **0.22 (0.01)** |  |

In addition, the cross-validation data is used to quantify the prediction performance. For each data set, all three error measures as well as the prediction uncertainty are computed (see Section 2.4). First, the relationship between prediction error and uncertainty is visualized in Fig. 2. This demonstrates that the smallest errors go hand in hand with the lowest uncertainty and, conversely, that a high certainty implies a small error. Furthermore, despite substantial uncertainty in the predictions for the macaque cerebral cortex in particular (middle panel of Fig. 2), most of the errors have a magnitude below one. This indicates that even when the predictions are uncertain, they are at most one category away from the true connection weight. Second, Table 2 shows the error measures for each of the different connectomes. These results indicate that the latent space model with dimensionality optimized through cross-validation consistently outperforms both baselines.

In the remainder of this section, each of the predicted connectomes is considered in detail.

### Predicting the macaque visual system

In Fig. 3A, the tracer data for the macaque visual system is shown, together with the expected posterior predicted connectome (i.e. the mean of the posterior samples). The predictions are based on a 2D latent space. Figure 3B shows the posterior distributions of the elements of **f**_*ij*_, i.e. the probability of an absent or present connection. The results are separated into those for the observed connections and those for unknown connections. For the observed connections, we see that the posterior distributions closely match the observed value (as indicated by the dotted line). This shows that the model is able to capture the observed structure well. For the unobserved connections, the results demonstrate that these are not simply copies of the distributions that correspond to the observations. For absent connections, the mean of the distribution is lower (i.e. an absent connection is less likely), and there is more uncertainty in the distribution as shown by its larger width. It follows that for present connections the mean is larger (i.e. a present connection is more likely). Indeed, we observe that the predicted connectome is slightly denser than the observed connections on their own, with a mean density of 48.4% (SD = 0.02), compared to an empirical density of 0.44. Figure 3C shows for each of the possible connections the width of the largest credible interval. The credible intervals range from 0 to 1, indicating that for some connections the model is entirely certain about its prediction, while for other connections the model cannot decide whether a connection should be absent or present. The structure of the upper panel of Fig. 3C shows that the largest uncertainty is, unsurprisingly, for the unobserved connections, but at the same time the model is confident about the prediction for other unknown connections, as can be seen in the lower panel of Fig. 3C. A list of the predicted connections in descending order of certainty is provided in D. Note that a higher latent dimensionality could decrease the prediction uncertainty, but this comes at the cost of losing generalizability; less uncertainty would lead to more prediction error.

**Figure 2.**
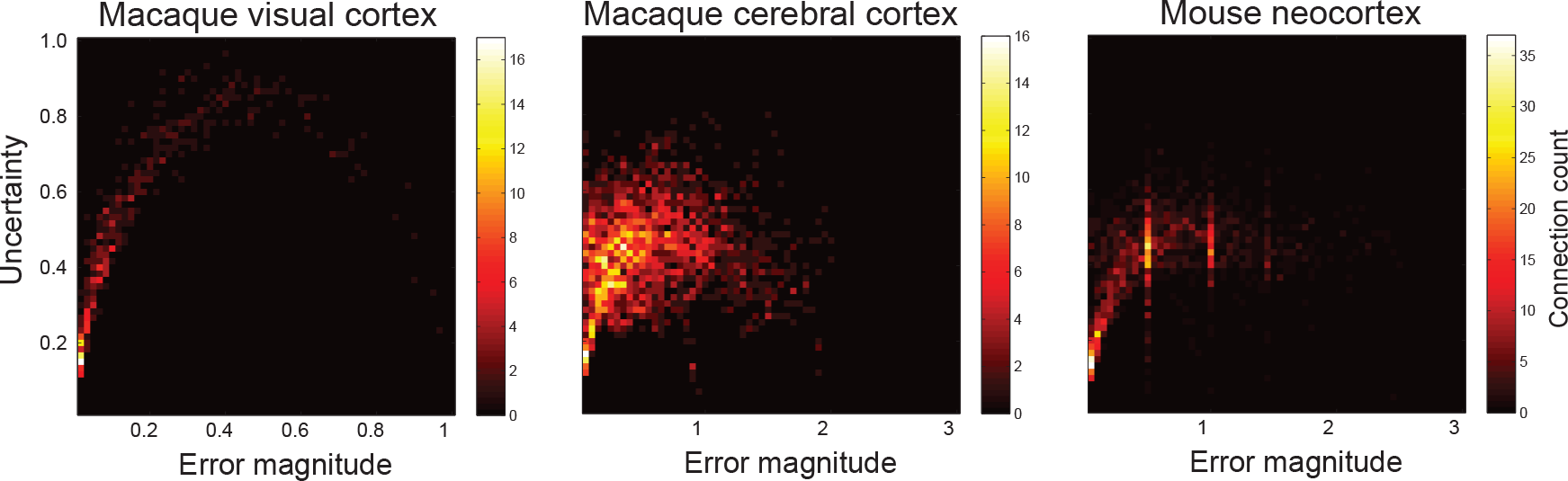
Prediction error and uncertainty. The relationship between prediction error (the absolute difference between the posterior expectation and the true value of the connection) and uncertainty (the width of the associated credible interval of the prediction). Colors are determined by the number of connections that lie within each cell; warmer colors indicate more connections. Note that for the mouse neocortex data, the prediction error is averaged over errors with the anterograde and retrograde data.

**Table 3.** Regions with a high (relative) increase in degree, based on the predicted connections for the macaque visual system. Shown are the top ten regions with the largest difference in fraction of observed connections 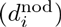 and fraction of predicted connections 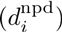. Scores are normalized according to the number of observed or total number of connections, respectively (see text).

| Outgoing |  |  |  |  |  | Incoming |  |  |  |  |  |
| --- | --- | --- | --- | --- | --- | --- | --- | --- | --- | --- | --- |
| Region | $d_i^{\text{nod}}$ | $d_i^{\text{npd}}$ | Region | $d_i^{\text{nod}}$ | $d_i^{\text{npd}}$ | Region | $d_i^{\text{nod}}$ | $d_i^{\text{npd}}$ | Region | $d_i^{\text{nod}}$ | $d_i^{\text{npd}}$ |
| AITd | 0.24 | 0.45 | DP | 0.40 | 0.55 | CITd | 0.06 | 0.32 | LIP | 0.47 | 0.65 |
| PITv | 0.21 | 0.42 | PITd | 0.25 | 0.39 | PITd | 0.13 | 0.39 | VP | 0.35 | 0.52 |
| PO | 0.39 | 0.58 | 7a | 0.45 | 0.58 | VOT | 0.50 | 0.71 | MIP | 0.00 | 0.16 |
| CITd | 0.24 | 0.39 | MSTl | 0.42 | 0.55 | PIP | 0.37 | 0.55 | MDP | 0.00 | 0.16 |
| CITv | 0.24 | 0.39 | STPp | 0.36 | 0.48 | V2 | 0.38 | 0.55 | V3a | 0.44 | 0.55 |

The predicted additional connections are not distributed homogeneously across the network. Instead, some nodes with only a few observed connections are predicted to become highly interconnecting hubs. Table 3 lists the ten regions with the most salient difference 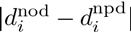, when comparing the observed connections and the predicted connectome, distinguishing outgoing and incoming connectivity. For example, the outgoing connectivity of the dorsal anterior inferotemporal cortex (AITd) increases substantially, as out of the ten unknown outgoing connections, 6.96 (SD=0.83) are predicted to be present.

Clustering the visual system connectome resulted in a division into occipitoparietal and parietotemporal areas, as shown in Fig. 4. Both the latent space prediction as well as the baseline model result in a division into two clusters. However, the latent space connectome results in a more modular division, as shown by the modularity score citepLeicht2008 of 0.25 compared to 0.20 in the baseline. However, the actual cluster assignments have changed slightly as well. As can be seen from Fig. 4, the medial intraparietal area, the medial dorsal parietal area and area 7a have been assigned differently between the two approaches.

**Figure 3.**
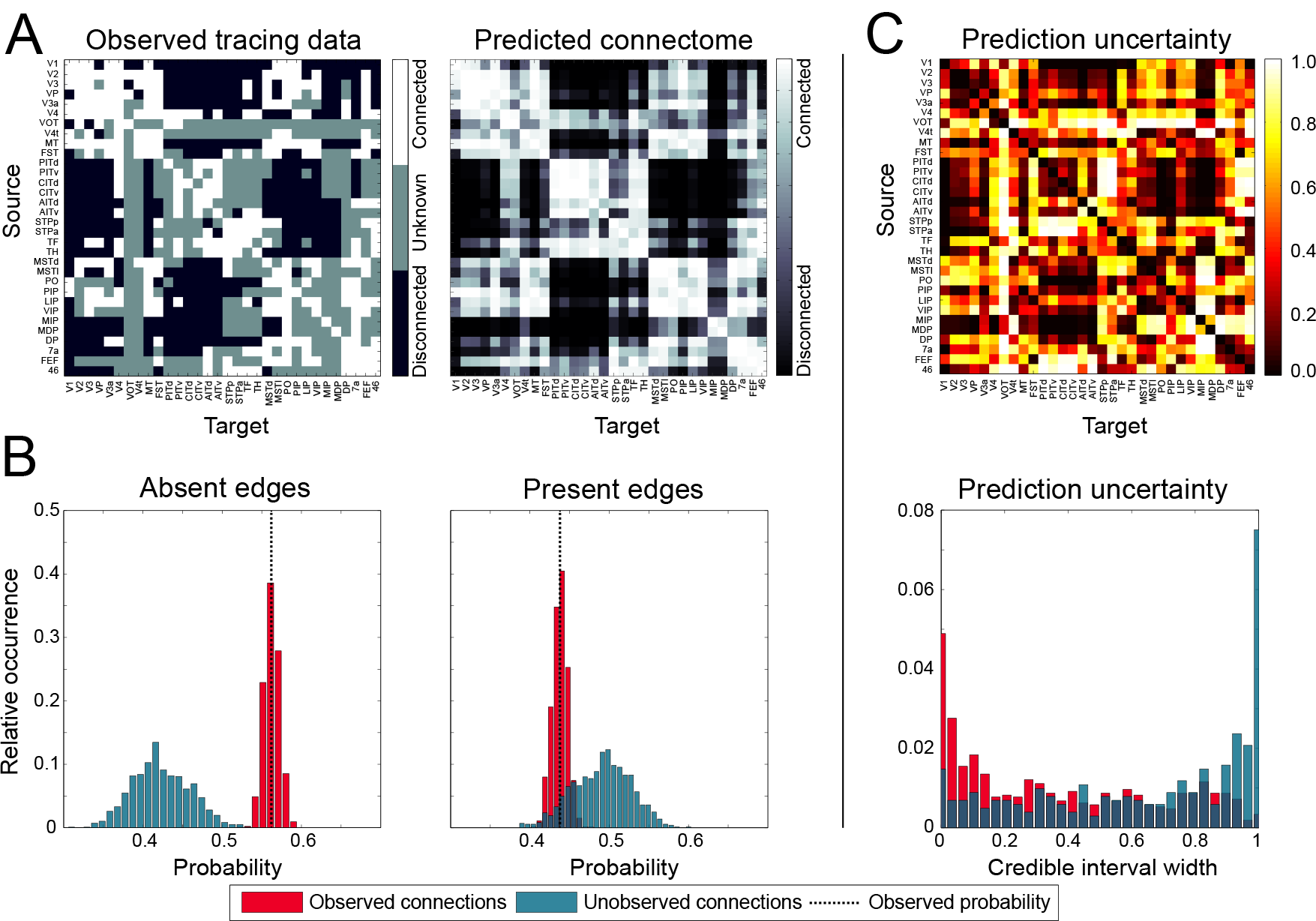
Macaque visual system connectivity. **A**. The observed tracing data for the macaque visual system (left) (Felleman & Van Essen 1991), the corresponding predicted connectome (right), based on the 2D latent space model. **B**. The predicted fraction of absent edges (left panel) and present (right panel), for observed and unobserved connections. For comparison, the dotted line shows the fraction of present edges in the observed connections. **C**. The uncertainty associated with each of the predicted connections.

### 3.2. Predicting the macaque cerebral cortex

While the visual system data contains sporadic missing data, for the macaque cerebral cortex (Markov et al. 2014) the majority of the connections are actually unobserved. The data is not missing at random, but systematically; retrograde tracers have been injected into 29 areas, which implies that bidirectional connectivity is only known for the subset of nodes within these 29 areas, and that retrograde connections are only known from any of the 91 areas to one of the areas of the subset (see the left panel of Fig. 5A). Despite this large amount of missing data, the latent space model may be used to predict the edge weights of the missing connections. As shown in Table 2, the macaque cerebral cortex network was best embedded into a 1D space. A consequence of this low-dimensional latent space is that the model prevents overfitting on the few observed connections. The predicted connectome is shown in the right panel of Fig. 5A. Figure 5B shows the proportions of each connection class (i.e. ‘absent’, ‘sparse’, ‘moderate’ and ‘strong’) for the predicted connectome, for either observed or unobserved connections, as well as the empirically observed frequency of each class. The figure demonstrates that, similar to the visual system data, the connections as predicted by the latent space model closely resemble the empirical edge class distribution, with slightly fewer absent connections and slightly more moderate and strong connections.

Figure 5C shows the uncertainty associated with the predictions. As to be expected, the uncertainty is highest for the unobserved connections, and lowest for those connections for which both anterograde and retrograde data was provided (i.e. the subnetwork of 29 regions). The widths of the credible intervals range from 0 to 0.93. A list of the predicted connections in descending order of certainty is provided in D.

**Figure 4.**
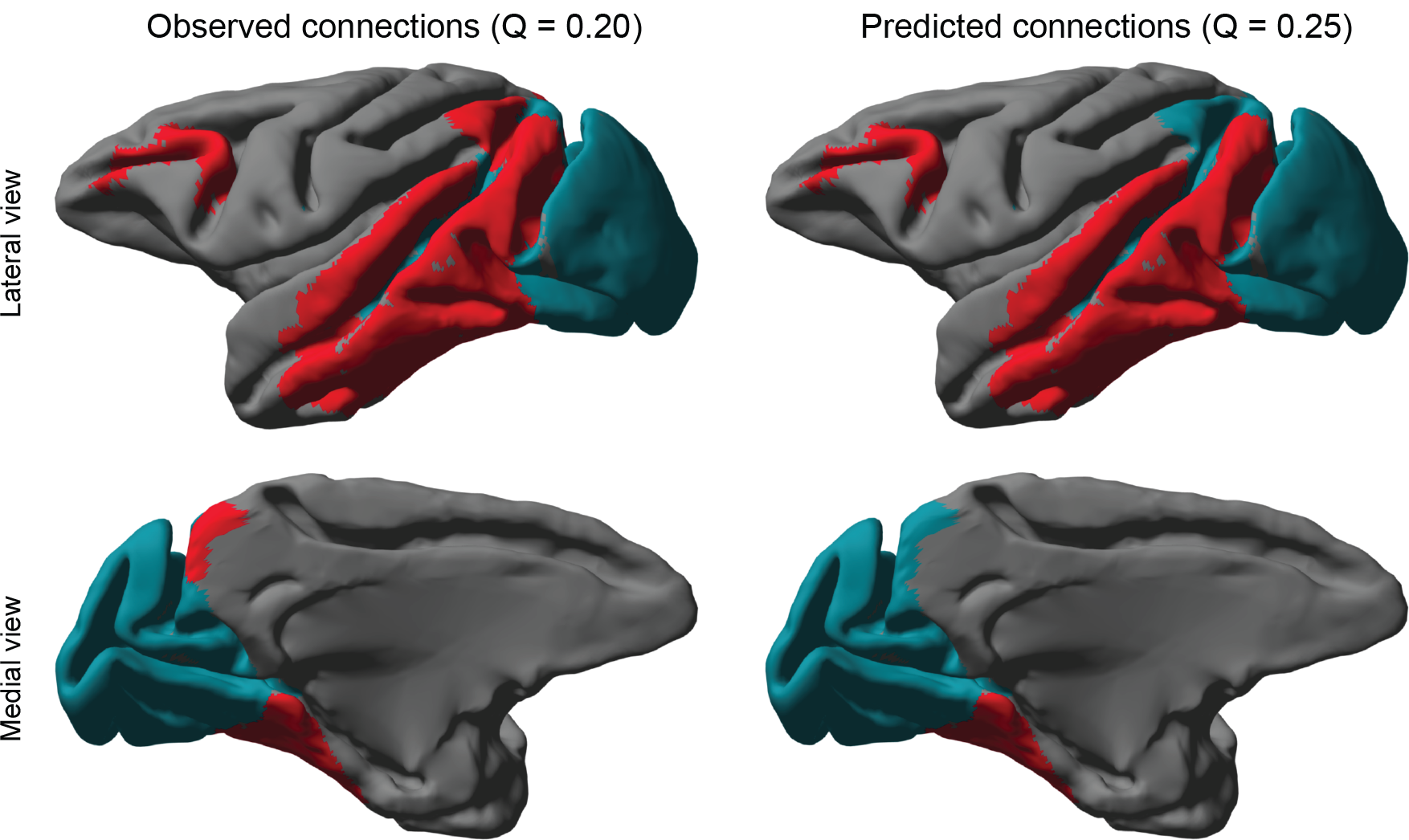
Clustering of the predicted macaque visual connectome. The figures on the left show the clustered visual system using the baseline model (see main text), while the figures on the right display the resulting clusters when connections have been predicted by the 2D latent space model. Both approaches result in a clustering of two clusters.

**Table 4.** Regions with the largest change in (relative) degree based on the predicted connections for the macaque cerebral cortex. Shown are the top twenty regions with the largest difference in fraction of observed connections 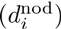 and fraction of predicted connections 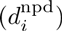. Scores are normalized according to the number of observed or total number of connections, respectively (see text). Note that the degree scores represent the average connection weight and range from 0 (absent) to 3 (strong).

| Incoming |  |  |  |  |  |  |  |  |  |  |  |
| --- | --- | --- | --- | --- | --- | --- | --- | --- | --- | --- | --- |
| Region | $d_i^{\text{nod}}$ | $d_i^{\text{npd}}$ | Region | $d_i^{\text{nod}}$ | $d_i^{\text{npd}}$ | Region | $d_i^{\text{nod}}$ | $d_i^{\text{npd}}$ | Region | $d_i^{\text{nod}}$ | $d_i^{\text{npd}}$ |
| PIR | 0.28 | 0.56 | 25 | 0.48 | 0.70 | Pi | 0.86 | 1.03 | STPc | 1.76 | 1.92 |
| V6 | 0.45 | 0.72 | V6A | 0.90 | 1.11 | 8m | 1.59 | 1.75 | 46d | 1.52 | 1.68 |
| SUB | 0.34 | 0.61 | PBr | 1.07 | 1.28 | TEO | 0.90 | 1.06 | ENTO | 1.10 | 1.26 |
| Core | 0.76 | 0.99 | 10 | 1.00 | 1.20 | STPr | 1.59 | 1.75 | 9/46d | 1.52 | 1.67 |
| Pro.St. | 0.55 | 0.77 | LB | 1.14 | 1.33 | TEpd | 1.10 | 1.27 | 8B | 1.55 | 1.71 |

In contrast to the macaque visual system, the predicted connections for the macaque cerebral cortex data follow the distribution of connectivity found in the data more closely. Table 4 lists for the regions with the largest difference in (weighted) incoming degree, when comparing the predicted connectome with the observed data. The differences in mean degree are comparatively small when considering that individual connections range on a scale from 1 (absent connection) to 4 (strong connection).

The resulting predictions were clustered and compared to the baseline model. This time the predicted connectome could be clustered into two clusters with a corresponding modularity score of 0.15. The baseline model instead resulted in four clusters, with a modularity score of 0.11, indicating that the predicted connectome resulted in a more modular network. Figure 6 shows the clusterings projected onto the cortical surface, which indicate that the difference in clustering is mostly due to the merger of frontal and parietal cortex, as well as a merger between temporal and occipital cortex.

**Figure 5.**
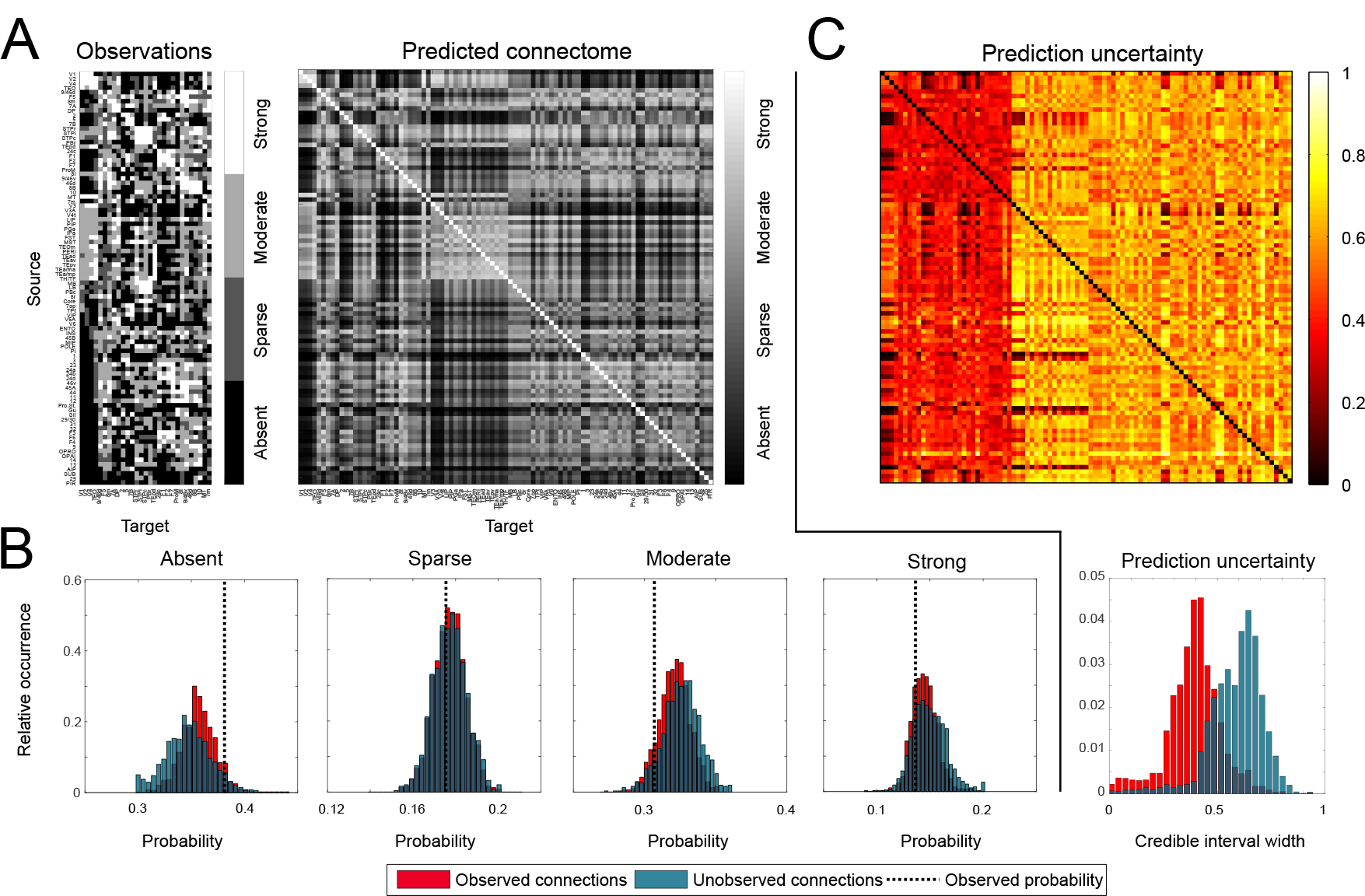
CMacaque cerebral cortex connectivity. **A**. The observed retrograde tracing results for the macaque cerebral cortex (Markov et al. 2014) (left) and the mean predicted connectome for all 91 regions using a 2D latent space (right). **B**. The distribution of connection weights for all connections on the prediction, only those that were previously unobserved and the observed relative frequencies of each class. **C**. The uncertainty associated with the predictions, for each possible connection (top) and as a histogram (bottom).

### 3.3. Integrating anterograde and retrograde data

Instead of predicting unobserved connections, the latent space model may also be used to integrate different modalities. Here, we combine both anterograde and retrograde tracing data collected for the mouse neocortex (Zingg et al. 2014) into a unifying estimate of the underlying connectome. Despite the aforementioned experimental variability, we expect most of the connections to be reciprocal. The model captures the remaining asymmetry using the directional effects parameters. As shown in Table 2, the best generalization performance is obtained using a 2D latent space when using either retrograde or anterograde tracers on their own. However, once the data are combined, a 4D space is optimal instead.

The first row of panels in Fig. 7A shows the predicted connectome for the mouse neocortex using either a single data source with a latent space dimensionality of two, or the combined data sources with a latent space dimensionality of four. The overall structure of the matrices appears to be the same in either setting. The histograms in Fig. 7B indicate that the latent space for the 4D data fusion model attempts to find consensus between the two modalities; the distributions of connection classes lie between the two observed values. This explains the need for a higher number of dimensions than previously encountered: the higher dimensionality allows the model to conform to both (contradicting) sources of data by forming a sort of ‘average’ connectome.

**Figure 6.**
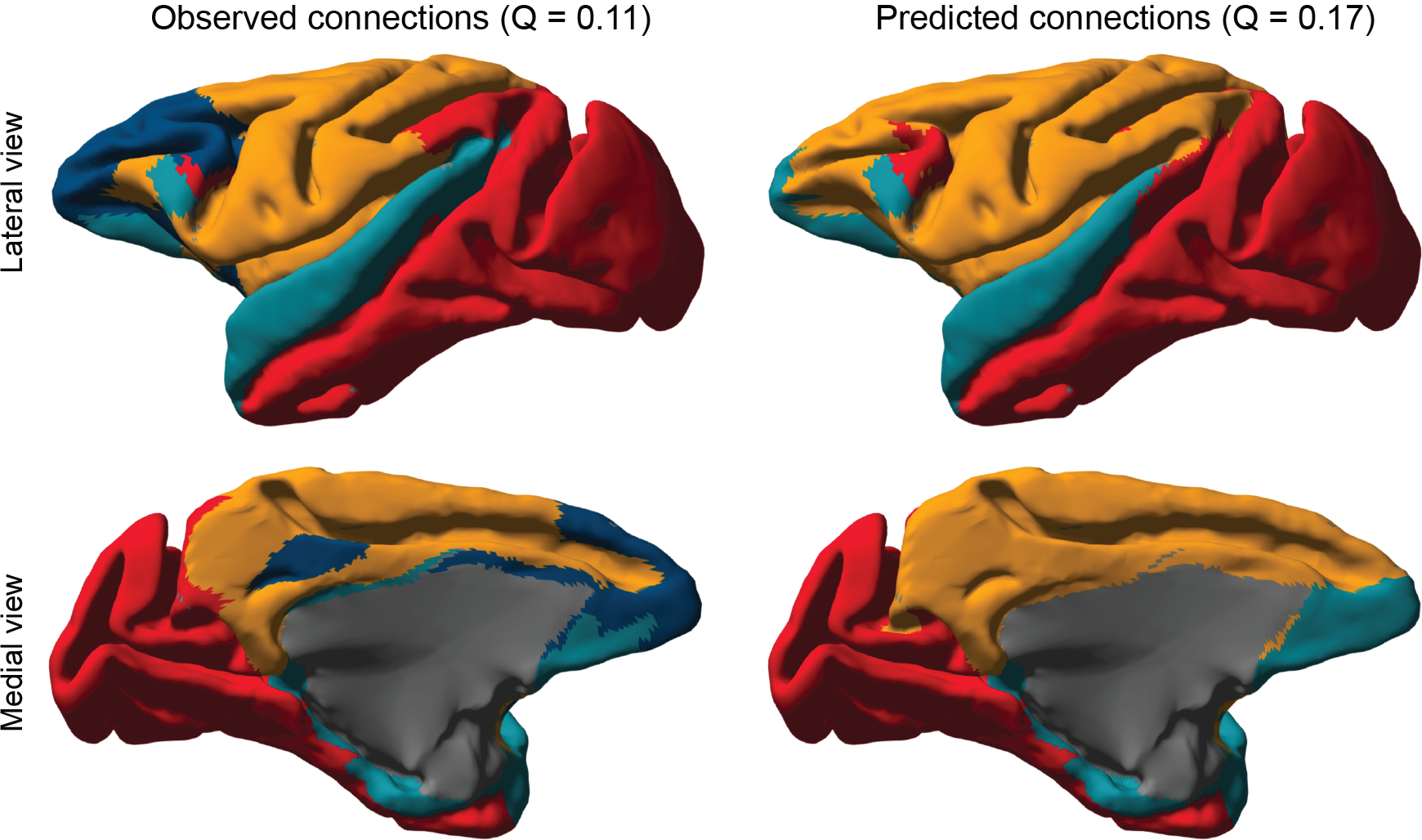
Clustering of the predicted macaque cerebral connectome. The figures on the left show the clustered cortex using the baseline model (see text), while the figures on the right display the resulting clusters when connections have been predicted by the 1D latent space model.

This intuition is confirmed by the edge-wise correlation between the mean of the anterograde and retrograde predictions and the predictions using data fusion: *ρ* = 0.97, *p* < 0.001, compared to a correlation between the anterograde and retrograde predictions of *ρ* = 0.93, *p* < 0.001. In other words, the top-rightmost adjacency matrix in Fig. 7A is approximately the mean of the other two. However, the prediction certainty increases by using both sources of data, as can be seen in Fig. 7B. When using just the anterograde data, the average uncertainty over all the connections is 0.35 (SD = 0.19), for just the retrograde data this is 0.32 (SD = 0.20) and when both tracer results are combined the average uncertainty is 0.29 (SD = 0.18). This demonstrates that although the predicted connectome using data fusion is approximately the mean of the predicted connectomes for either data set, data fusion provides the benefit of decreasing the uncertainty in the spatial embedding and consequently the connection prediction.

## 4. Discussion

We have demonstrated that connectivity can be predicted between regions for which no data has been observed directly. We accomplished this by estimating a latent space in which the nodes of the connectome are embedded. In this latent space, the distances between nodes determine the probability or strength of the corresponding connections: nodes that are close together are more likely to be connected (or have a higher connection strength) than nodes that are far apart. The latent space model (LSM) was applied to predict connectivity for two connectomes of the macaque, as well as to integrate anterograde and retrograde tracer data for the mouse neocortex. The predictions were accompanied by low average errors, outperforming the two baseline approaches for link prediction in which either the connection class distributions were used, or a zero-dimensional latent space that is only able to predict on the basis of the directional effects (i.e. node degrees). These findings demonstrate that the spatial embedding approach captures important features of connectivity.

**Figure 7.**
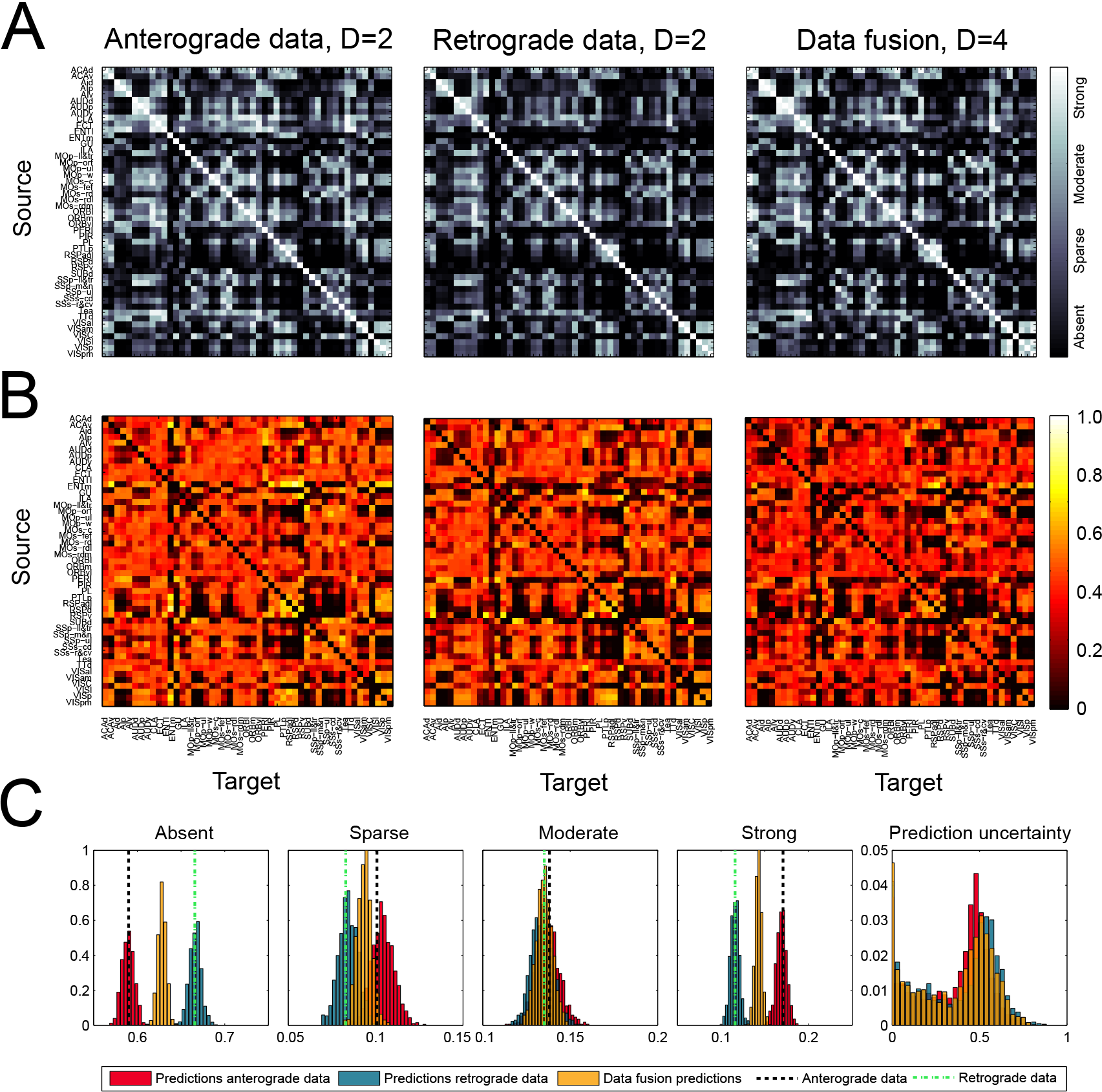
Combined mouse neocortex connectivity. **A**. Predicted connectivity for the mouse neocortex (Zingg et al. 2014), using either anterograde data only (using two latent dimensions), retrograde data only (using two latent dimensions), or the combination of both (using four latent dimensions, see main text). **B**. The distribution of the edge classes in the predicted connectome, compared to the values in the observed data.

When many of the connections are actually not observed (as for the macaque cerebral cortex connectome), the model may be uncertain about its predictions. As the described LSM is probabilistic, this prediction uncertainty can be made explicit. This revealed that even though on average the model performs well, depending on the data a substantial number of (potential) connections could not be predicted with confidence. This is due to too few data points being available for the involved regions, so that the LSM cannot fixate their positions in the latent space. The representation of uncertainty provides target areas for additional experiments with novel tracers. Those connections with maximal uncertainty should be probed first in future work, so that these connections become known, and the latent space parameters with the most degrees of freedom become anchored, which will propagate to making other predictions more certain as well. Our approach is therefore applicable both for prediction of unseen connections, as well as for guidance of optimal experimental design (Priebe et al. 2013, Pukelsheim 1993).

The choice for a latent space model for link prediction in brain connectivity is motivated by findings that show the probability of two regions being connected (and the strength of this connection) is inversely correlated with the Euclidean distance between them (Bullmore & Sporns 2012, Ercsey-Ravasz et al. 2013, Raj & Chen 2011, Chen & Hall 2006). This indicates that indeed the wiring of the brain is constrained by its geometry. Remarkably, a small number of latent dimensions appeared to suffice in order to model each of the considered connectomes. Only in the mouse neocortex data, a 4D latent space resulted in optimal generalizability instead. We speculate that this is due variability between individual mice and layer-specific connectivity, which creates differences between the anterograde and retrograde observations.

There are a number of ways in which the model may be extended in future work. First, following (Durante & Dunson 2014, Miller et al. 2009), a nonparametric variant of the model may be constructed that learns the dimensionality D of the latent space on-the-fly. This avoids the need to split the available data into train and test sets, as all the data can be used to train the model and learn *D* simultaneously. Another extension could describe the FLNe weights in the macaque cerebral cortex data (Markov et al. 2014) as a continuous variable rather than the currently used thresholded categorization. We have refrained from this approach in order to present one model that was applicable to all three data sets, but it is to be expected that continuous weights better inform the latent distances than the ad-hoc ordinal representation of connection strengths, and subsequently increase prediction performance.

A few studies are related to the presented work and have analyzed the spatial embeddedness of structural connectivity. For example, Ercsey-Ravasz et al. (2013) use the 29 × 29 submatrix of fully observed connectivity in the macaque cerebral cortex to predict global graph-theoretical properties of the full 91 × 91 connectome. Here, it is assumed that the fully observed submatrix is representative of the entire connectome. This relates to our observation about the (relative) degree of regions in the data and in the predicted connectome, which have been found to be quite similar in the macaque cerebral cortex data. Bassett et al. (2010) consider a more abstract notion of topological dimensionality. Here, different nervous systems as well as computer circuits are studied and found to have a higher topological complexity than the physical embedding of the networks would suggest. As Bassett et al. (2010) argue, this implies that connectomes optimize for a trade-off between minimal wiring length and maximum topological complexity. Finally, Song et al. (2014) demonstrate that many large-scale features of connectivity in mouse, macaque and human can be explained by simple generative mechanisms based on spatial embedding.

We have demonstrated the usage of the latent space model for prediction of connectivity on animal tracer data. The advantage of tracer data over other modalities is their reliability (as, for example, dMRI-based structural connectivity estimates are often accompanied by uncertain estimates (Hinne et al. 2013, Janssen et al. 2014)), which allowed us to evaluate the performance of link prediction using the latent space model. In terms of application however, other modalities may benefit more from our approach. For example, dMRI in combination with tractography is often used to estimate structural connectivity *in vivo*, making it applicable to human subjects. This approach has a number of well-known shortcomings that affect the resulting connectomes (Jones et al. 2013, Schultz et al. 2014, Reveley et al. 2015). By using the LSM approach, connections that are difficult to estimate in living human subjects may be predicted from the connections that were more easily obtained. Furthermore, recent advancements in electron microscopy imaging have enabled connectivity analysis at the single-cell resolution (Helmstaedter 2013, Lichtman et al. 2014). Here, link prediction may be used to complete the connectivity that has not yet been probed and at the same time to obtain insight in the possible spatial embedding of connectivity at this scale.

## Appendix A. Generative model for ordinal latent space

The formal description of the generative model for the latent space embedding with asymmetric effects is as follows:

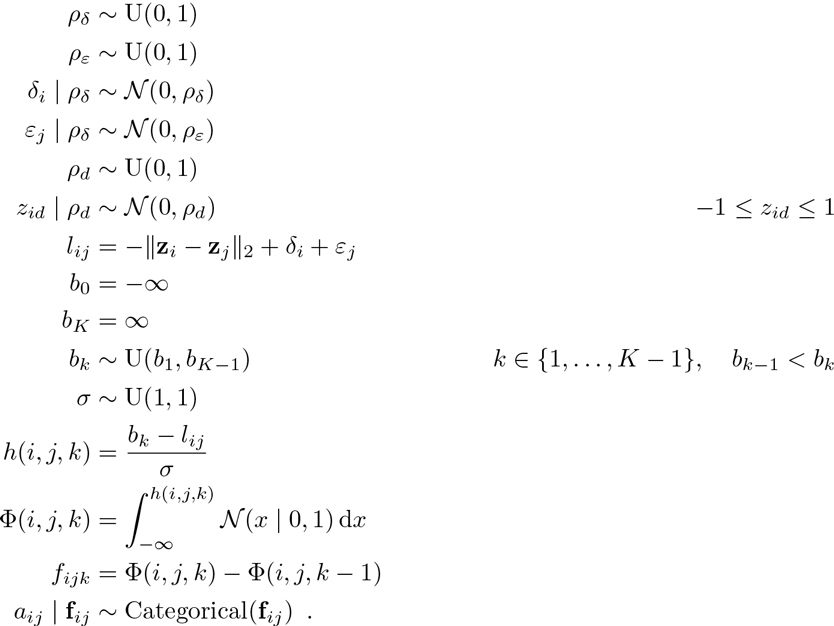

Here, *a*_*ij*_ represents the categorical class of the connection between nodes *i* and *j*. Note that the symbol ~ should be read as ‘follows the distribution’. As the latent space model considers only the relative distances between nodes, the positions **z** may be arbitrarily scaled and translated throughout the latent space. In order to have the posterior distribution be consistent across different samples, we constrain the scale by setting 0 < σ ≤ 1 and we constrain the positions to lie within the *D*-dimensional unit hypercube by requiring −1 ≤ *z*_*id*_ ≤ 1. Extending the model to integrate both anterograde and retrograde tracing data is straightforward by adding the likelihood term

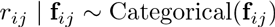

to the model. Here, *r*_*ij*_ represents the retrograde connection while the original *a*_*ij*_ parameter represents the anterograde connection. Notably, both types of observations depend on the same latent distances.

## Appendix B. Approximate inference and scalability

The model is implemented in MatlabStan (Lau 2015), which provides an interface between Matlab and the Stan probabilistic programming language. This software implements the no-U-turn Hamiltonian Monte Carlo sampler (Hoffman & Gelman 2014). For each different model, four parallel sampling chains are executed. Convergence to the posterior distribution is determined by computing the potential scale reduction factor (PSRF) (Gelman & Rubin 1992) for parameters *l*_*ij*_, **f**_*ij*_ and σ. Once all PSRF scores are below 1.1 (typically after 6 000–10000 iterations), the chains are considered to be converged. Subsequently, the chains are merged and downsampled to 1 000 samples for efficient further analysis.

Per iteration of the Hamiltonian Monte Carlo algorithm, a total of (*D* + 2)*p* + *K* + 2 parameters need to be estimated. However, making general claims about the computational cost of the HMC approach is difficult as convergence depends the ease of which the latent positions can be determined, which in turn depends on the dimensionality *D* and the latent structure in the data. For example, we noticed that during the cross-validation procedure, the *D* = 1 case was easy to compute, but difficult to obtain convergence for (taking as much as 10000 iterations), while higher dimensional latent spaces had more computational cost per iteration, yet converged much faster (in as few as 4 000 iterations). To provide a guideline for the efficiency of the approach, Fig. 8 shows the computation time per 100 iterations for the data used in this paper, using a single Intel Xeon CPU E5-2670 @ 2.60GHz per sampling chain. The approximately linear trends that are shown in these results may be used to extrapolate running times for connectomes with a larger number of nodes than used here. For example, prediction of connectivity for a connectome of 1 000 nodes, using a 2-dimensional latent space, should take roughly four days to compute, using the same hardware.

**Figure 8.**
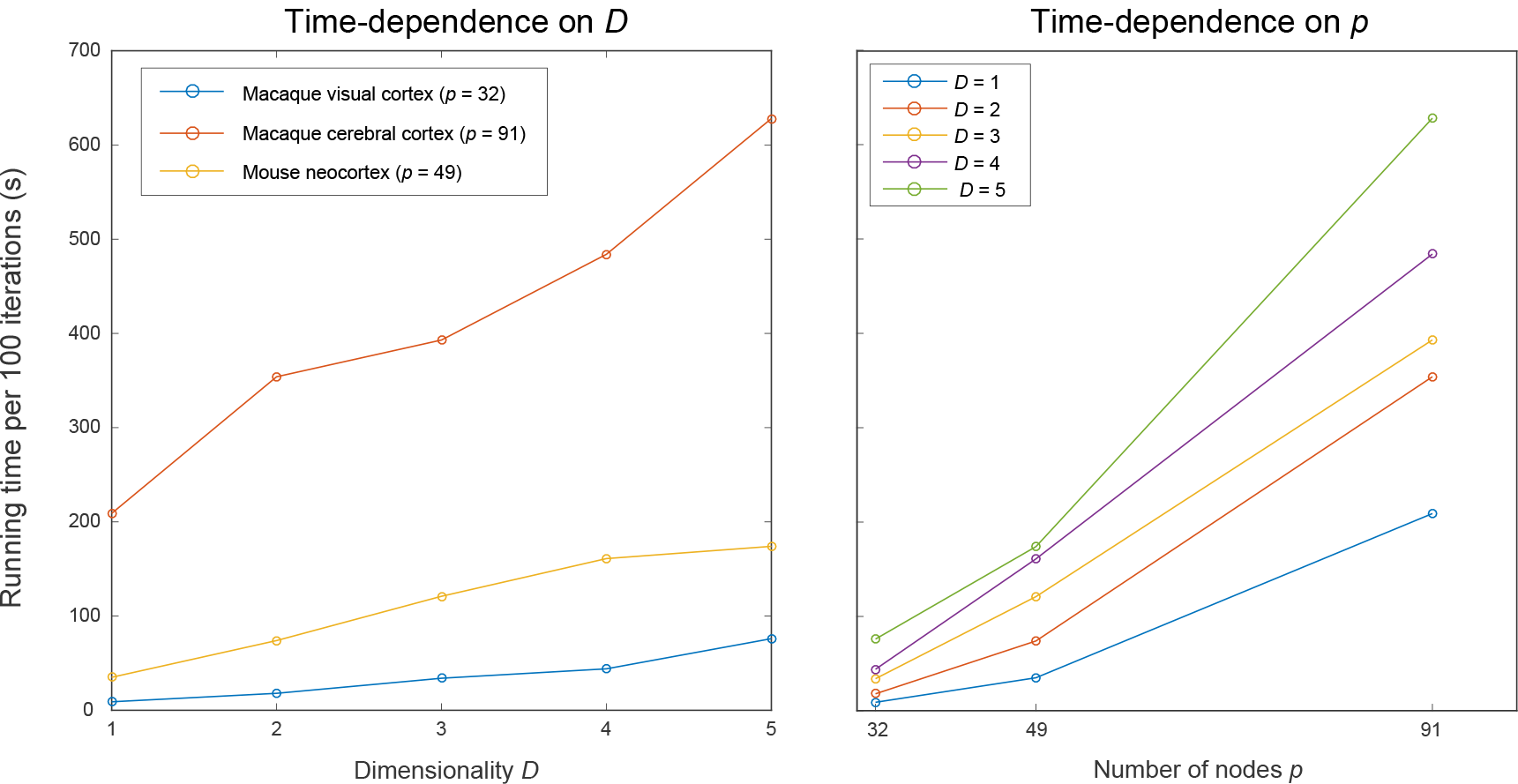
Running time of 100 iterations of the Hamiltonian Monte Carlo algorithm on each of the different data sets that are considered in the main text. The figures confirm the (approximately) linear dependence on the most important parameters of the model; the dimensionality D (left panel) and the number of nodes in the connectome p (right panel).

## Appendix C. Predictions for the different dimensionalities

The predictions using different dimensionalities (in the range [0,…, 6]) are shown in Fig. 9 for the macaque visual system, in Fig. 10 for the macaque cerebral cortex and in Fig. 11 for the mouse neocortex.

**Figure 9.**
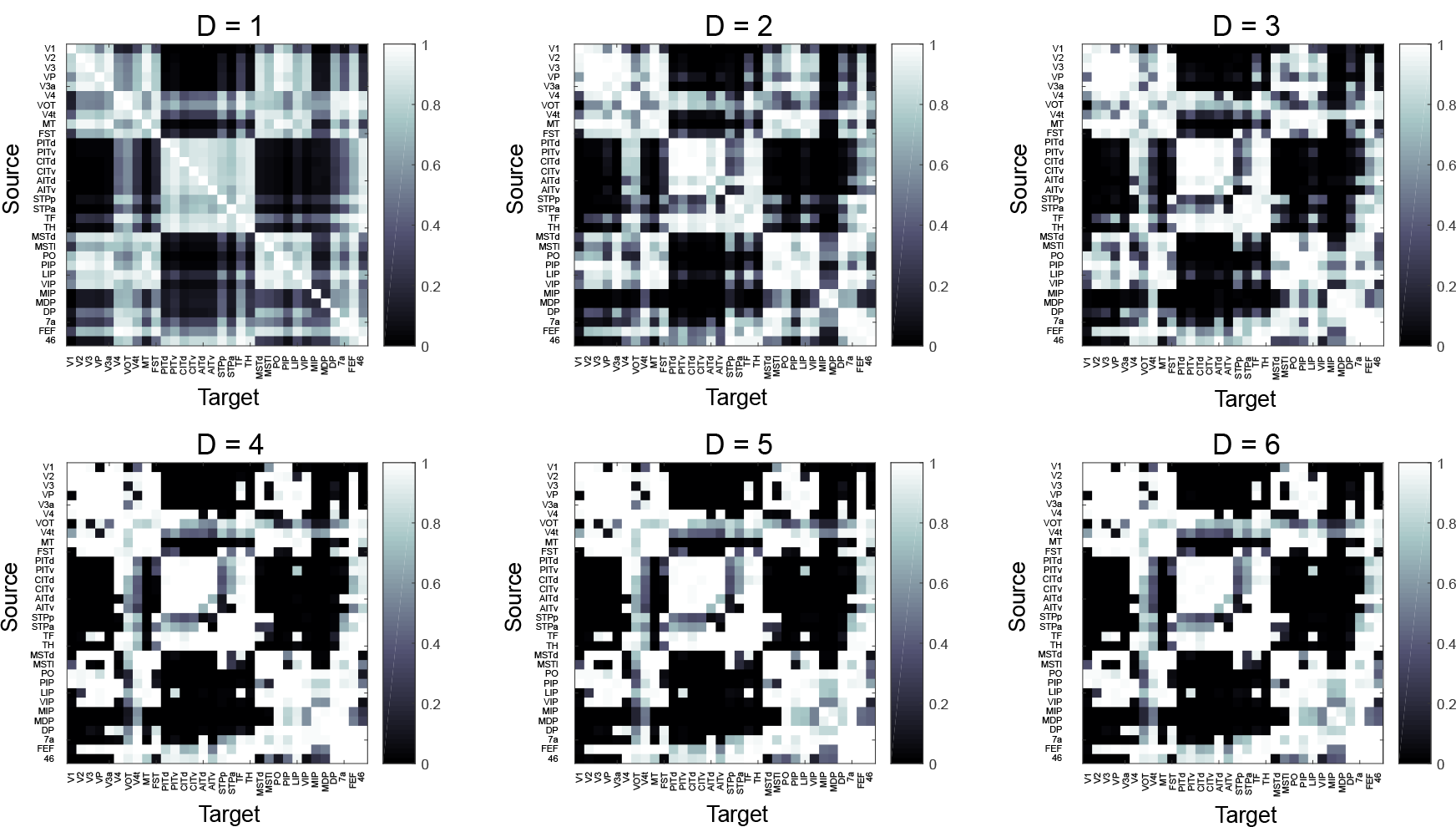
Predictions for the macaque visual system. Posterior expectations of the connectivity matrix **A** for different numbers of latent dimensions.

**Figure 10.**
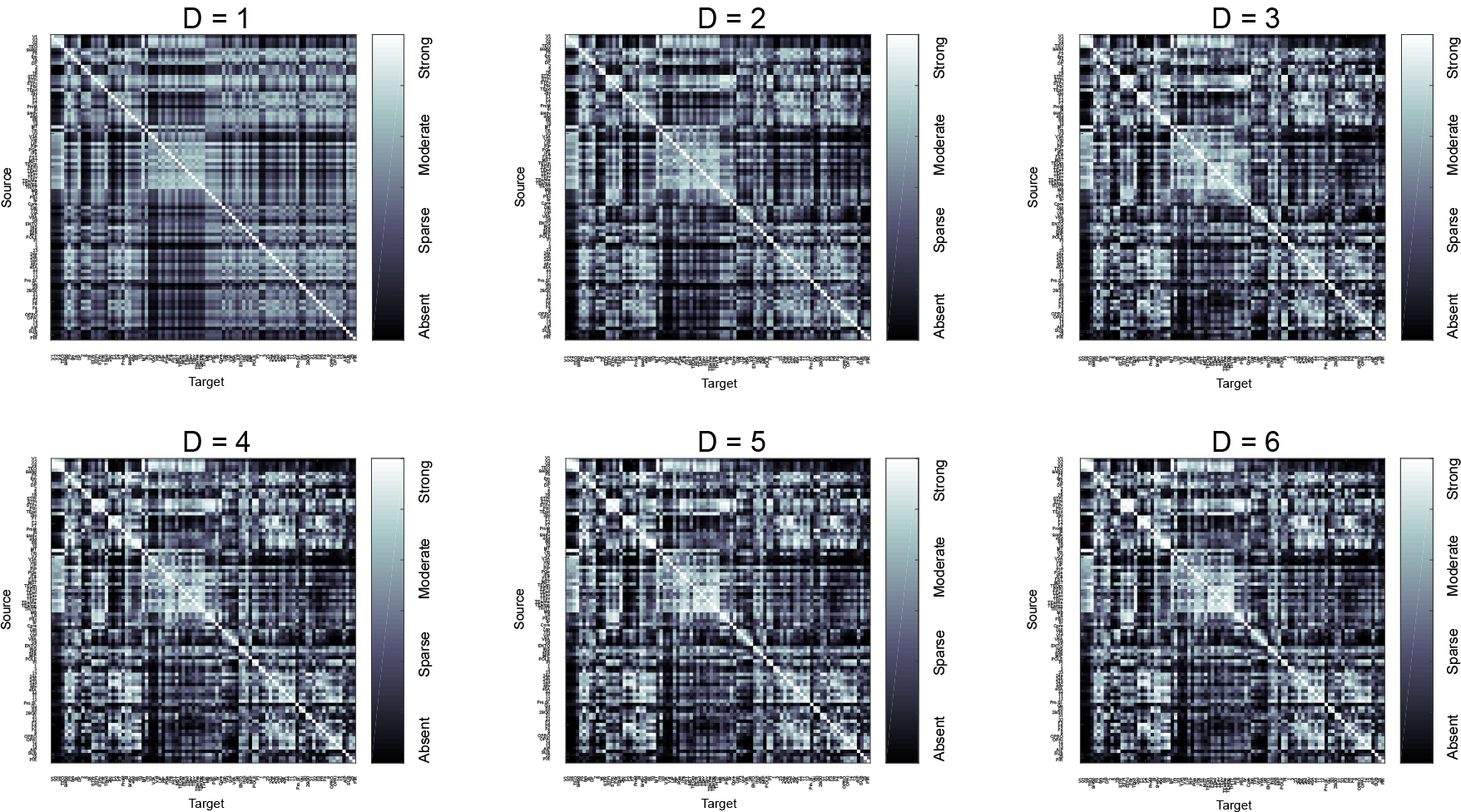
Predictions for the macaque cerebral cortex. Posterior expectations of the connectivity matrix **A** for different numbers of latent dimensions.

**Figure 11.**
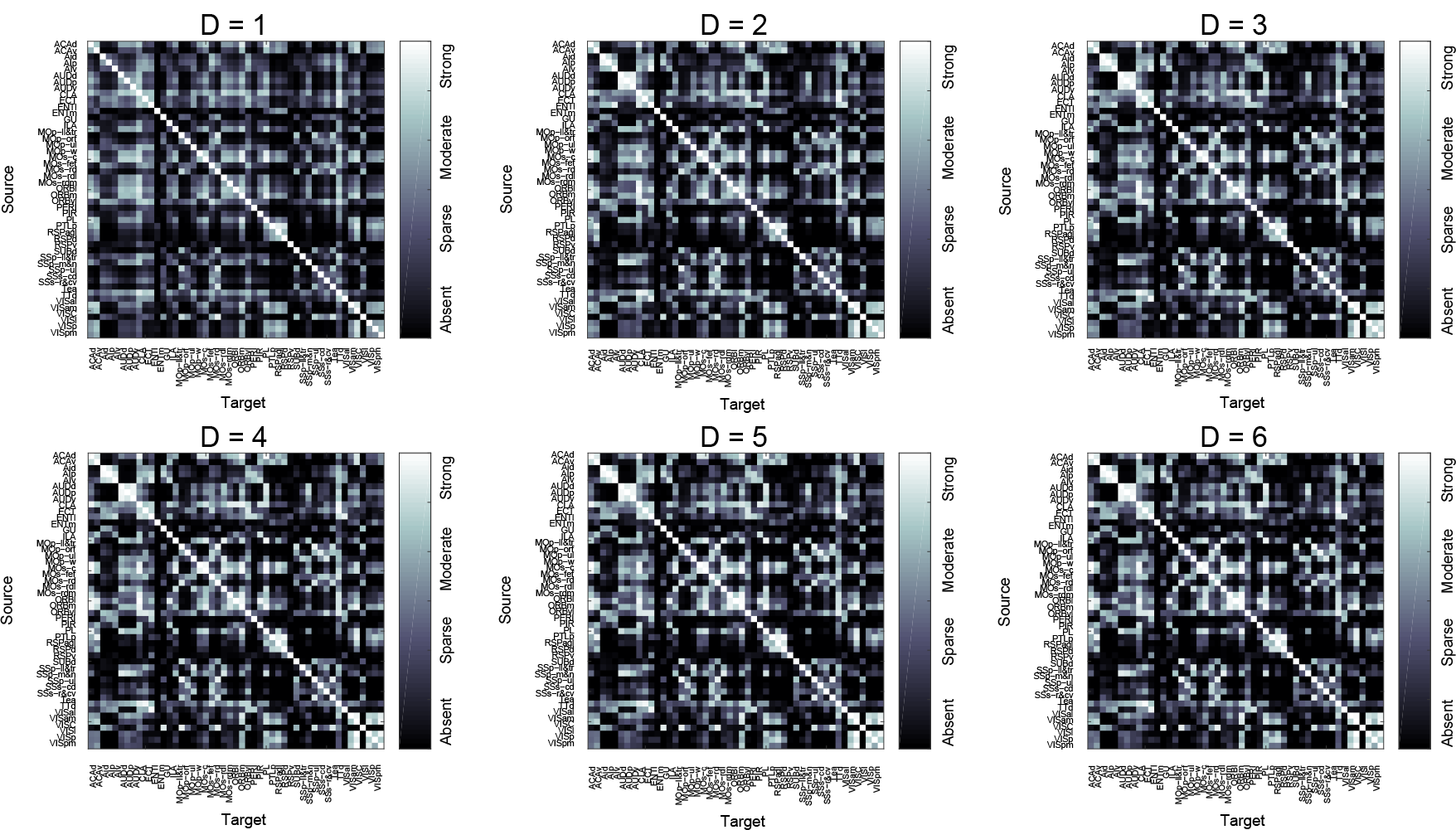
Predictions for the mouse neocortex. Posterior expectations of the connectivity matrix **A** for different numbers of latent dimensions.

## Appendix D. Predicted connections

The top forty predicted connections, ordered by their prediction certainty (i.e. smallest associated credible interval) for both the macaque visual cortex and macaque cerebral cortex are shown in Table 5 and Table 6, respectively. The full listings can be found in Supplementary Information 1 and 2. For the full region labels we refer the reader to Supplementary Information 3.

**Table 5.** The top forty predicted unknown connections for the macaque visual cortex, sorted by decreasing uncertainty. The predicted connection strength is shown as *â*_*ij*_, while *c*_*ij*_ indicates the width of the associated credible interval.

| # | source | target | $\hat{a}_{ij}$ | $c_{ij}$ | # | source | target | $\hat{a}_{ij}$ | $c_{ij}$ |
| --- | --- | --- | --- | --- | --- | --- | --- | --- | --- |
| 1 | CITd | MDP | 0.00 | 0.00 | 21 | MDP | TF | 0.01 | 0.06 |
| 2 | CITv | MDP | 0.00 | 0.00 | 22 | TH | TF | 0.99 | 0.06 |
| 3 | PITd | MDP | 0.00 | 0.00 | 23 | TF | FEF | 0.99 | 0.07 |
| 4 | CITd | MIP | 0.00 | 0.00 | 24 | LIP | V4t | 0.99 | 0.08 |
| 5 | CITv | MIP | 0.00 | 0.00 | 25 | AITv | DP | 0.01 | 0.09 |
| 6 | PITv | MDP | 0.00 | 0.00 | 26 | V4t | VIP | 0.99 | 0.09 |
| 7 | PITd | MIP | 0.00 | 0.00 | 27 | STPp | 7a | 0.99 | 0.09 |
| 8 | PITv | MIP | 0.00 | 0.00 | 28 | FEF | TF | 0.99 | 0.09 |
| 9 | PO | PITd | 0.00 | 0.00 | 29 | MIP | TF | 0.01 | 0.09 |
| 10 | PIP | V3A | 1.00 | 0.01 | 30 | VIP | V4t | 0.99 | 0.10 |
| 11 | MSTl | DP | 1.00 | 0.02 | 31 | PITd | PITv | 0.99 | 0.10 |
| 12 | MSTd | MSTl | 1.00 | 0.02 | 32 | V3 | VP | 0.99 | 0.10 |
| 13 | V3A | PIP | 1.00 | 0.03 | 33 | CITd | CITv | 0.99 | 0.11 |
| 14 | AITd | TH | 1.00 | 0.03 | 34 | VIP | DP | 0.99 | 0.11 |
| 15 | PIP | V2 | 1.00 | 0.03 | 35 | V4t | DP | 0.99 | 0.12 |
| 16 | V4t | MSTd | 1.00 | 0.03 | 36 | 46 | PIP | 0.02 | 0.12 |
| 17 | MSTl | MSTd | 1.00 | 0.03 | 37 | 7a | STPp | 0.99 | 0.13 |
| 18 | MDP | TH | 0.00 | 0.03 | 38 | V4t | LIP | 0.99 | 0.13 |
| 19 | TF | TH | 1.00 | 0.05 | 39 | CITd | DP | 0.02 | 0.14 |
| 20 | MIP | TH | 0.01 | 0.06 | 40 | CITv | CITd | 0.99 | 0.15 |

**Table 6.** The top forty predicted unknown connections for the macaque cerebral cortex, sorted by decreasing uncertainty. The predicted connection strength is shown as *â*_*ij*_, while *c*_*ij*_ indicates the width of the associated credible interval.

| # | source | target | $\hat{a}_{ij}$ | $c_{ij}$ | # | source | target | $\hat{a}_{ij}$ | $c_{ij}$ |
| --- | --- | --- | --- | --- | --- | --- | --- | --- | --- |
| 1 | V1 | Gu | 0.00 | 0.00 | 21 | V1 | 3 | 0.00 | 0.04 |
| 2 | V6 | Piriform | 0.00 | 0.01 | 22 | Piriform | MIP | 0.00 | 0.04 |
| 3 | Subiculum | 1 | 0.00 | 0.01 | 23 | V4t | S2 | 0.01 | 0.04 |
| 4 | V4t | Gu | 0.00 | 0.01 | 24 | Gu | TEOm | 0.01 | 0.04 |
| 5 | Gu | V4t | 0.00 | 0.01 | 25 | V3 | Gu | 0.01 | 0.04 |
| 6 | Gu | V3A | 0.00 | 0.02 | 26 | V1 | F4 | 0.01 | 0.05 |
| 7 | 1 | Subiculum | 0.00 | 0.02 | 27 | Prostriate | AIP | 0.00 | 0.05 |
| 8 | Prostriate | 1 | 0.00 | 0.02 | 28 | MIP | Piriform | 0.01 | 0.05 |
| 9 | Gu | V3 | 0.00 | 0.02 | 29 | Gu | PIP | 0.01 | 0.05 |
| 10 | V3A | Gu | 0.00 | 0.02 | 30 | PIP | Gu | 0.01 | 0.05 |
| 11 | Subiculum | AIP | 0.00 | 0.02 | 31 | 1 | TEad | 0.01 | 0.05 |
| 12 | V6 | 25 | 0.00 | 0.03 | 32 | 1 | Piriform | 0.01 | 0.05 |
| 13 | Piriform | V6 | 0.00 | 0.03 | 33 | V3A | OPAI | 0.01 | 0.05 |
| 14 | V1 | S2 | 0.00 | 0.03 | 34 | V4t | 3 | 0.01 | 0.05 |
| 15 | V1 | OPAI | 0.00 | 0.03 | 35 | V1 | 1 | 0.01 | 0.05 |
| 16 | TEad | 1 | 0.00 | 0.04 | 36 | Piriform | 1 | 0.01 | 0.06 |
| 17 | 1 | Prostriate | 0.00 | 0.04 | 37 | Subiculum | MIP | 0.01 | 0.06 |
| 18 | V6 | OPAI | 0.00 | 0.04 | 38 | ProM | V4t | 0.01 | 0.06 |
| 19 | Piriform | VIP | 0.00 | 0.04 | 39 | Subiculum | VIP | 0.01 | 0.06 |
| 20 | AIP | Subiculum | 0.00 | 0.04 | 40 | TEad | AIP | 0.01 | 0.06 |

